# Fast Parallel Algorithm for Large Fractal Kinetic Models with Diffusion

**DOI:** 10.1101/275248

**Authors:** A. A. Popov, S.-C. Lee, P. P. Kuksa, J. D. Glickson, A. A. Shestovb

## Abstract

Chemical kinetic simulations are usually based on the law of mass action that applies to behavior of particles in solution. Molecular interactions in a crowded medium as in a cell, however, are not easily described by such conventional mathematical treatment. Fractal kinetics is emerging as a novel method for simulating kinetic reactions in such an environment. To date, there has not been a fast, efficient, and, more importantly, parallel algorithm for such computations. Here, we present an algorithm with several novel features for simulating large (with respect to size and time scale) fractal kinetic models. We applied the fractal kinetic technique and our algorithm to a canonical substrate-enzyme model with explicit phase-separation in the product, and achieved a speed-up of up to 8 times over previous results with reasonably tight bounds on the accuracy of the simulation. We anticipate that this technique and algorithm will have important applications to simulation of intra-cell biochemical reactions with complex dynamic behavior.

## INTRODUCTION

The kinetics of a crowded medium does not behave like that of a freely diffusing solution [1]. The rate of reaction in such a medium varies with time and space, and is, therefore, not constant, as opposed to conventional kinetics [2, 3, 4]. Classical Michaelis-Menten kinetics [5] is, therefore, not applicable to all cases [6]. The effective fractal dimension (the dimension of an average particle diffusion trajectory) of a crowded medium is often closer to 2 and not 3 [7], and, thus, the assumptions made by the law of mass action [8, 9] are violated. The addition of explicit particle-particle phase-separation is also inconsistent with traditional kinetic particle simulation [10, 11, 12], and forces us to look at methods in which this type of complex individual particle diffusion is modeled explicitly. Attempts to model phase-separation in terms of reaction-diffusion equations [13, 14] are all limited in application and do not correspond well to the behavior of individual molecules in the cell.

Fractal kinetics represents one method of resolving these issues with conventional simulations. Assuming a finite lattice, we define the cells of the lattice to be all its primitive cells and the size of the lattice to be the number of cells that it contains. We define the neighbors of a cell to include all the near neighbors. For a 2D square lattice, those are the cells that share an edge with the cell, and for a 3D cubic lattice those that share a face with the cell. We additionally define a metric on the cells: the distance between two cells will be the length of the shortest near neighbor path between the cells, such that the neighbors of a cell will have distance 1 from it, their neighbors (not already having been assigned a distance) to have distance 2, and so forth. For a 2D square lattice this is equivalent to the Manhattan distance.

In a fractal kinetic simulation, all particles are discrete and, thus, occupy one cell, or multiple ‘small enough’; particles occupy the same cell. No particle can occupy multiple cells. This will be the limiting factor of the spatial discretization. We define a particle to be any constituent in a kinetic equation, thus for the canonical substrate-enzyme reaction [9],

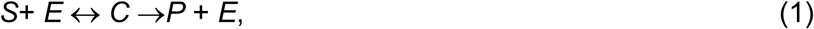

the reactants *S, E, C* and *P* are all particles. Via a method of iteration through the cells of the lattice, each cell is simulated individually, and can interact with itself and with any cells up to a specified distance, *d*_*i*_. An interaction may be a reaction with a neighbor, diffusion, change of state, breaking up into multiple particles, or anything else. Iterating through and simulating the same number of cells as the size of the lattice (including the simulation of empty cells) is a Monte-Carlo Step (MCS). For example, simulating the first cell of a 2 × 3 lattice five times and simulating the third cell once is a MCS.

With particle diffusion, creation of a safe parallel algorithm becomes much more difficult, as errors such as particle duplication (when two threads move the same particle into different cells) and particle extinguishing (when two threads move two particles into one cell) can exponentially harm the simulation. Even a single diffusion error could catastrophically alter the dynamics of the simulation. Our scheme, therefore, has to be completely devoid of diffusion errors. A large part of the difficulty in efficiently simulating fractal kinetics with diffusion stems from the choice of the specific way by which to iterate through the lattice cells. In order to simulate such a lattice one cannot simply simulate the cells sequentially, as that could potentially lead to bias in how certain particles diffuse. We will describe how both the previous naive algorithm, and our algorithm operate on an *L* × *L* square lattice, but both can be trivially generalized to any 2D or 3D lattice. By parallel, we mean that we have several threads, each operating on separate portions of the same data. For example, given two non-adjacent cells, it is possible to simulate both at the same time on separate threads. We define *T* to be the number of threads.

Previous algorithms attempting to model fractal kinetics [15, 16] opted for a naive approach, whereby, at each MCS, *L*^2^ lattice cells are chosen randomly (with the same cell can be picked multiple times), and are sequentially simulated. Whilst this does, on average guarantee that each cell will be simulated at approximately the same rate, it requires that there be an extremely fast way of procuring random numbers, as this problem is trivially on the order of *L*^2^. One of the fastest “good” random number generators available at the moment, xoroshiro128+, can generate one 64-bit number in about 0.87*ns* on a conventional consumer CPU, thus for a 1000 × 1000 lattice simulation running for 10^7^ MCS, we would spend about 10 days just on generating random numbers for lattice traversal. We are, thus, limited not only by the severe restriction of having one executing thread thereby not being able to utilize the full capabilities of modern multi-core machines, but also by the speed of our random number generator. Other lattice simulation parallelization schemes such as sub-lattice algorithms [17], are not built for simulations involving particle diffusion, and, thus, still require enormous amounts of random numbers, and can potentially introduce diffusion errors which could be rapidly magnified and dramatically alter the concentration of a quickly diffusing particle.

Analyzing the naive Monte-Carlo approach, as seen in Fig 1, we see that the simulated average number of cells fluctuates above and below the number of MCSs performed. On a large enough time scale, we expect any given cell to be---at some point in time---the lowest simulated cell, and the highest simulated cell. As our simulations involve particle diffusion, the more important metric is on the short time scale, since the cell can rapidly be simulated many times in a MCS or potentially not simulated at all for several MCSs. These data put severe limitations on the way a cell can be simulated. A sequential simulation via a method of permutation would not produce the same trajectories; thus any serious method must attempt to replicate this behavior.

**Fig 1.**
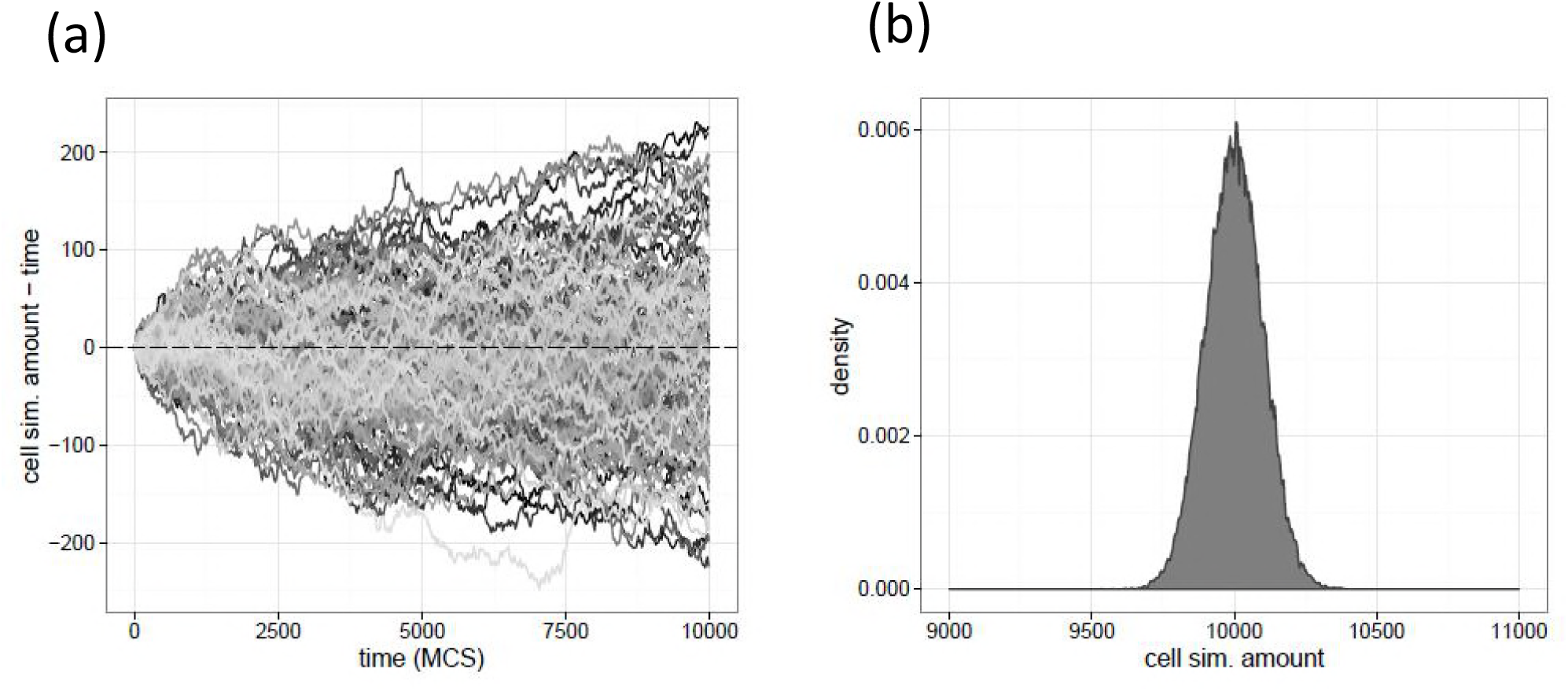
Naive algorithm analysis. Simulating an empty 200 × 200 (*L* = 200) lattice for 10^4^ MCS and counting the running total of the amount of cell simulations, yields the following two graphs. On the left is the relative cell simulation of the number of trajectories with respect to the MCS, showing a tendency of the individual cell simulation numbers to diverge. The end simulation numbers (right), however, all tend towards a Gaussian distribution.

## METHODS

To generalize the naive algorithm to finite-cell multi-threaded simulations, we introduce the concept of covers. We define a cover to be any set of sequences of cells in one lattice. For example, a set of two sequences, one traversing randomly the top half of a lattice, and one containing a length-one sequence of a cell in the bottom half would be a cover. The cover containing one random sequence of cells from the lattice of length *L* × *L* would be equivalent to one MCS of the naive algorithm. Thus, the naive algorithm can be thought of as such truly random cover at each MCS simulating the cells by its single sequence.

We extend the simulation of one sequence cover to multiple sequences. A cover having multiple sequences would simulate each of them in parallel in its own thread. To clarify, the sequence length would now be arbitrary, and both the lengths of the sequences and the respective cells in each would be chosen to fulfill some desired properties. We define the coverage of a cover to be a lattice containing counts of how many times a cell appears in all its sequences. The coverage of a collection of covers would be a per-cell sum of individual coverages. In the naive algorithm, the coverage of the cover for a single step would be a lattice containing integers from 0 to possibly (but not likely) *L*^*2*^, and the coverage of the whole algorithm would be a lattice whose elements would tend towards infinity with time, as each cover is independently drawn.

The foundation of our algorithm is based on the idea of generating covers such that parallel execution can safely (i.e. with no diffusion errors) take place on a single lattice. This is an example of the load balancing problem [18]. We, thus, try to eliminate both problems that we have identified previously. We will only rely heavily on generating random numbers for the initial formulation of the cover and will be able to operate safely in parallel on one lattice at a time without introducing any diffusion errors. To generate our covers, we first define one parameter, the maximal distance *d* = 2*d*_i_+1, as the distance that two cells simulated in parallel will not interact with the same cell at the same time. In conventional diffusion, where during one unit of simulation, the maximal event that can occur is a particle moving one cell distance, we have *d*_*I*_ = 1, and thus *d* = 3. We will use this parameter throughout the remainder of the calculations.

Consider an empty *L* × *L* square lattice and choose a random *L*^2^-length sequence of its cells with possible repetition and omission. We then operate on each cell in the sequence, one-by-one, either assigning it a thread, or leaving it unassigned. For each cell on which we operate, we take a closed neighborhood of cells around it with radius *d* –1, excluding itself. We then proceed with the following decision process:

- If the cell is already assigned to a thread, assign it to the same thread again.
- If there are no other assigned cells in the neighborhood, we assign the cell to a thread at random.
- If there are other assigned cells in the neighborhood, all assigned to one thread, we assign the cell to that thread.
- If there are cells such that two or more different threads are in the neighborhood, we do not assign the cell to any thread, and increment our collision counter.

When the collision ratio of the cover (collision counter divided by *L*^2^) becomes larger than our allowed collision ratio (*CR*), we stop. Splitting the processed sequence by assigning thread to a set of sequences generates a cover. Call the number of covers that we wish to generate *n*_*C*_. From the coverage of a given collection of *n*_*C*_ covers, we can generate a heat map in order to indicate which cells will be simulated more, and which will be simulated less. We ideally wish to generate a monotone heat map such that every cell is simulated the same number of times over our total collection of covers. As we can see from the heat map in Fig 2a, our initial algorithm is sub-optimal as the 2-3 cell wide perimeter has a slightly increased frequency of coverage and has the strongest coverage at the four corners. Also note the large difference between the maximally covered cell and the minimally covered cell.

**Fig 2.**
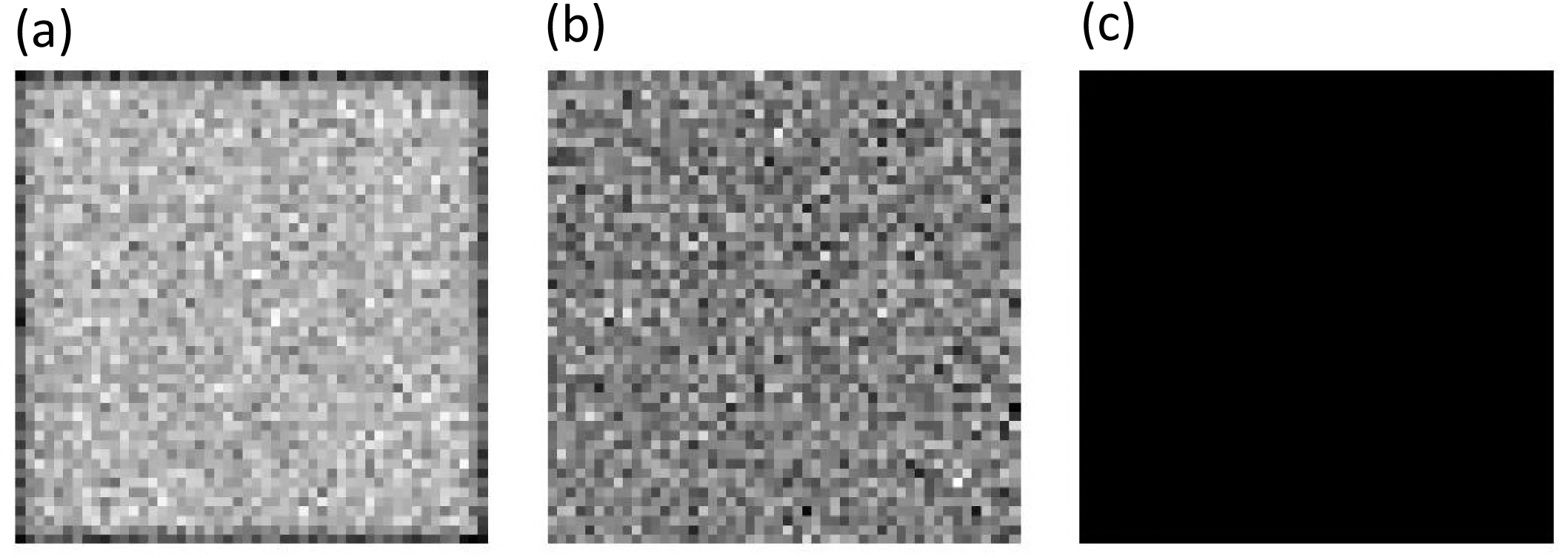
Heat map Analysis. (a) For *L* = 50, *n*_*C*_ = 25000, *CR* = 2^−6^, *T* = 4. Coverage min = 3434, coverage max = 4064. (b) For *L* = 50, *n*_*C*_ = 25000, *CR* = 2^−6^, *T* = 4 with cyclic boundaries. Coverage min = 3319, coverage max = 3765. (c) For *L* = 50, *n*_*C*_ = 25000, *CR* = 2^−6^, *T* = 4 with synthetic cyclic boundaries, and min-max correction. Coverage min = max = 3766. [22] Heat maps of several different ways of generating covers. The figure on the left represents a heat map of the coverage of the covers generated by our algorithm before any corrections. The figure in the middle is a heat map of the coverage of the collection of covers generated with synthetic cyclic boundary conditions imposed to alleviate the larger coverage on the boundary in the previous heat map. The figure on the right represents the heat map of the collection of covers generated by imposing both synthetic cyclic boundary conditions on the cover generation, and min-max cover padding. Full white represents the minimum coverage of the collection of covers, and full black represents the maximum coverage. Low *L*, high *n*_*C*_, and high *CR* are used for demonstrative purposes. The figure on the right is all black representing a uniform maximal coverage of the lattice.

The first type of correction is one that is immediately obvious. We introduce synthetic cyclic boundary conditions to correct the error on the perimeter. Such a correction leads us to generate a second heat map as seen in Fig 2b, which does not suffer on the edges. This, however, still leaves us with an imbalance in the maximal and minimally covered cells. The second type of correction is min-max correction. We take all the cells whose coverage is not maximal (e.g. for the covers of Fig 2b, we take all the cells such that their coverage is less than the maximal coverage of the set of covers, 3765) and generate a cover only over them, omitting all maximal points, and omitting duplication (to avoid generating an infinite number of covers). This procedure is repeated until the minimal and maximal coverages of the set of covers are equal. A monotone heat map resulting from this correction is depicted in Fig 2c. We are then left with a set of covers that uniformly covers the lattice, thereby producing the desired coverage effect. A pseudocode for this algorithm can be seen in Table 1. Figure 3 shows an example of a cover generated for two threads on a 50 × 50 lattice.

**Table 1.**
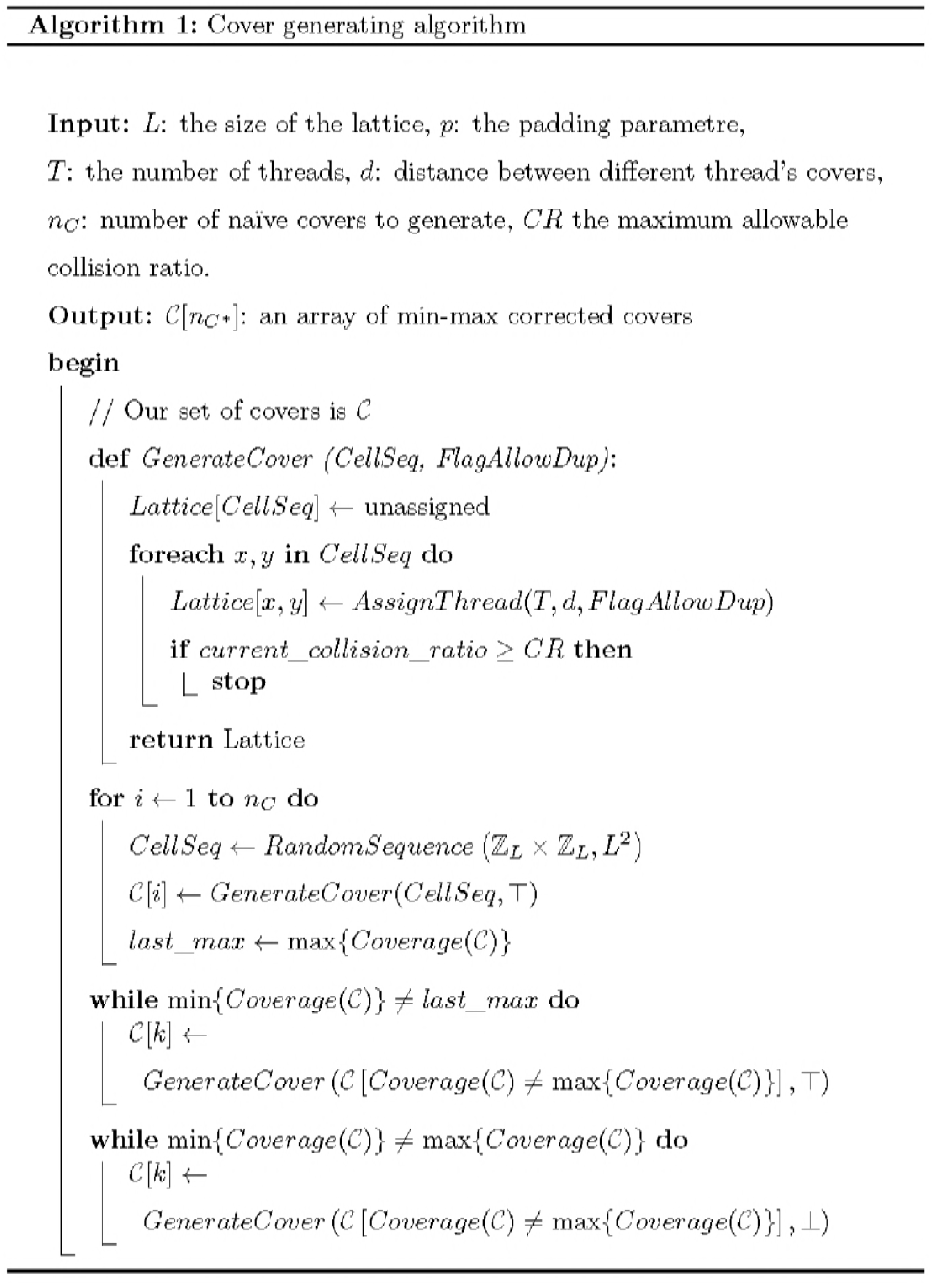

**Fig 3.**
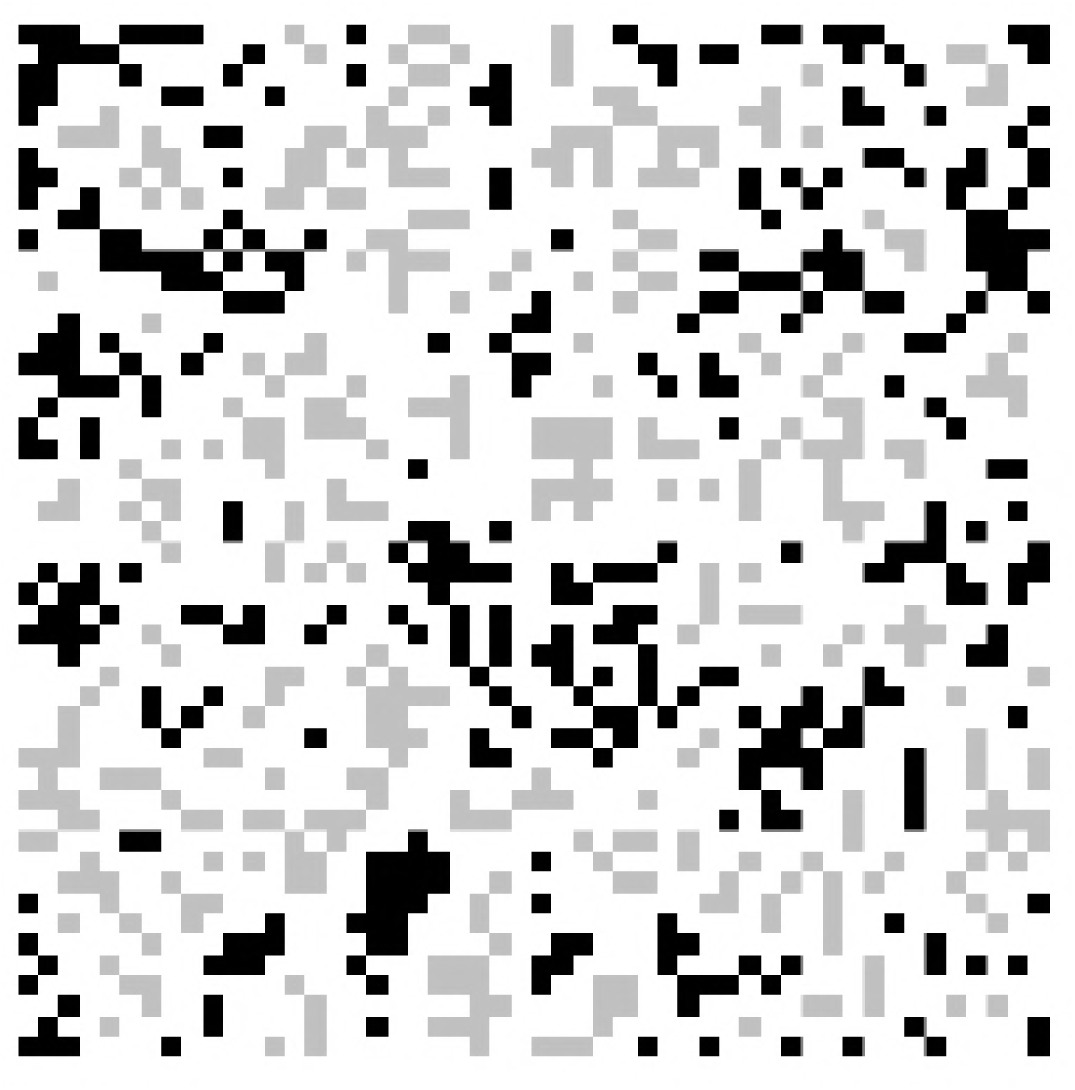
Sample Cover. A sample cover for a 50×50 (*L* = 50) lattice, with *T* = 2, *d*_*i*_ = 1, *d* = 3, *CR* = 2^−3^ and cyclic boundary conditions on cover generation. White represents no assigned thread, while the two shades of gray each represent a separate thread. The specific cell sequences are not visualized, but can be any random permutation of cells of the same color, and can include repeating cells. The minimum number of empty cells between any two cells of different colors is 2.

We first create *n*_*C*_ cyclic boundary covers for *T* threads over our *L* × *L* lattice. We then add additional covers for min-max corrections, obtaining *n*_*C\**_ covers. Having produced a set of covers that uniformly cover the lattice, we can subsequently define our fractal kinetic algorithm. Call Δ*MCS* the interval of MCS between our observations of the lattice, and let c be the coverage of a cover. For each Δ*MCS* we have to simulate *L*^*2*^Δ*MCS* cells. We create a sequence of covers at random (with repetition) up until a point where the sum of their respective coverages is greater than or equal to *L*^*2*^Δ*MCS*. The number of covers required to achieve this is *n*_*C*_^*n*^. Call 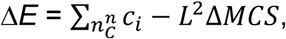 the number of extra cells that we simulate for this interval.

For the next interval, we subtract Δ*E* from the required coverage number, so that on average we simulate *L*^*2*^Δ*MCS* cells per observation interval. We then simulate the *n*_*c*_^*n*^ covers. Each subset of cells in a cover is simulated in parallel by a thread. As no two covers can be simulated at the same time---in order to avoid errors---we have to synchronize all threads after each cover simulation. Thread synchronization also introduces its own set of challenges; however, fast methods can mitigate many of these complications. A pseudocode for this is depicted in Table 2. Figure 4 shows a sample simulation with a sample cover whereby the fractal kinetic algorithm is executed over one cover in a safe and parallel way. Analyzing the algorithm that we have created, we see that our algorithm performs similarly to its naive counterpart, thus providing empirical evidence of its validity (Fig 5).

**Table 2.**
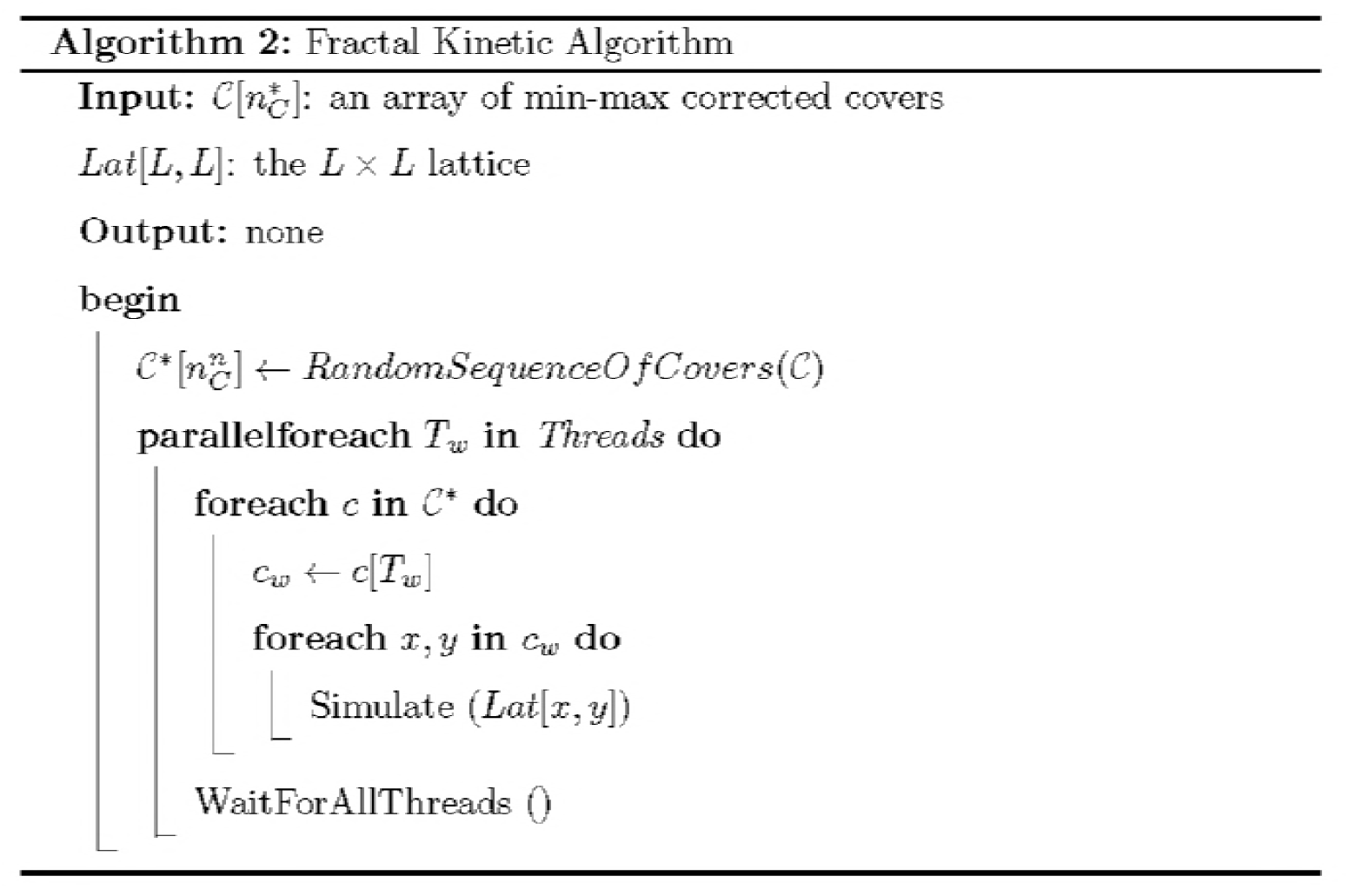

**Fig 4.**
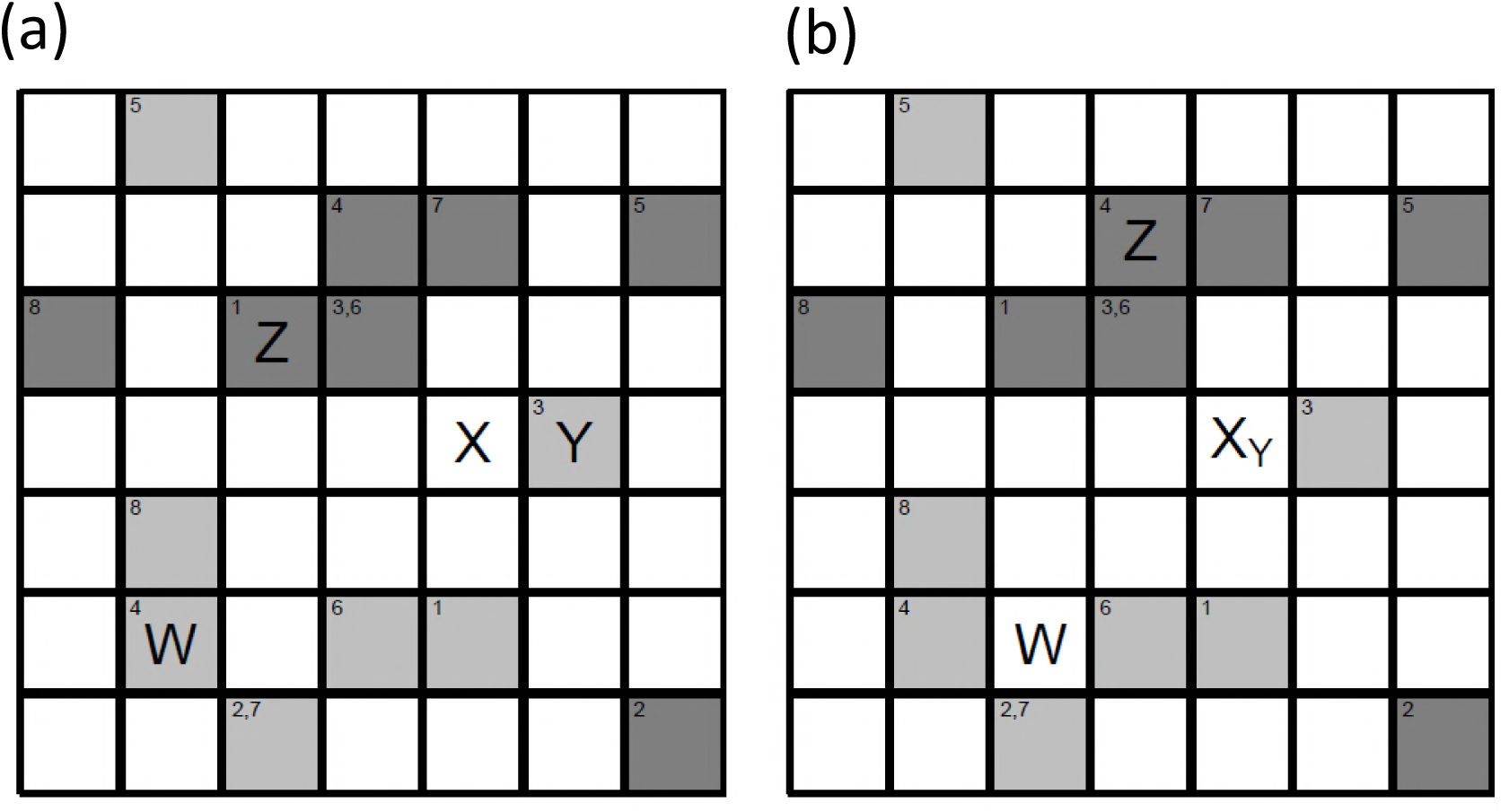
Example of the working algorithm for one sample cover. (a) The lattice configuration before the simulation of the cover. (b) The lattice configuration after the simulation of the cover. For *L* = 7, *T* = 2, a single cover (by hand) was constructed over the lattice. In the two graphics the two shades of gray represent a separate execution thread, and the numbers represent the order of execution in the thread (with some cells being executed multiple times). Particles *W, X, Y* and *Z* were inserted into the lattice simulating the cover. Particle *W* moved one unit to the right on the fourth step of the light-gray thread, and was then outside the scope of the cover. Particle *X* was outside the scope of the cover from the beginning, and, thus, did not move during the execution of the cover. Particle *Y* moved one unit to the left on the third step of the light-gray thread and interacted with particle *X* to create particle *X*_*Y*_, which was then outside the scope of the cover. Particle *Z* first moved to the right one unit on the first step of the dark-gray thread, then up one on the third step, then down one on the fourth step, and then up one again on the sixth step, ending its journey inside the scope of the cover, having moved four times and being two units away from its starting location.

**Fig 5.**
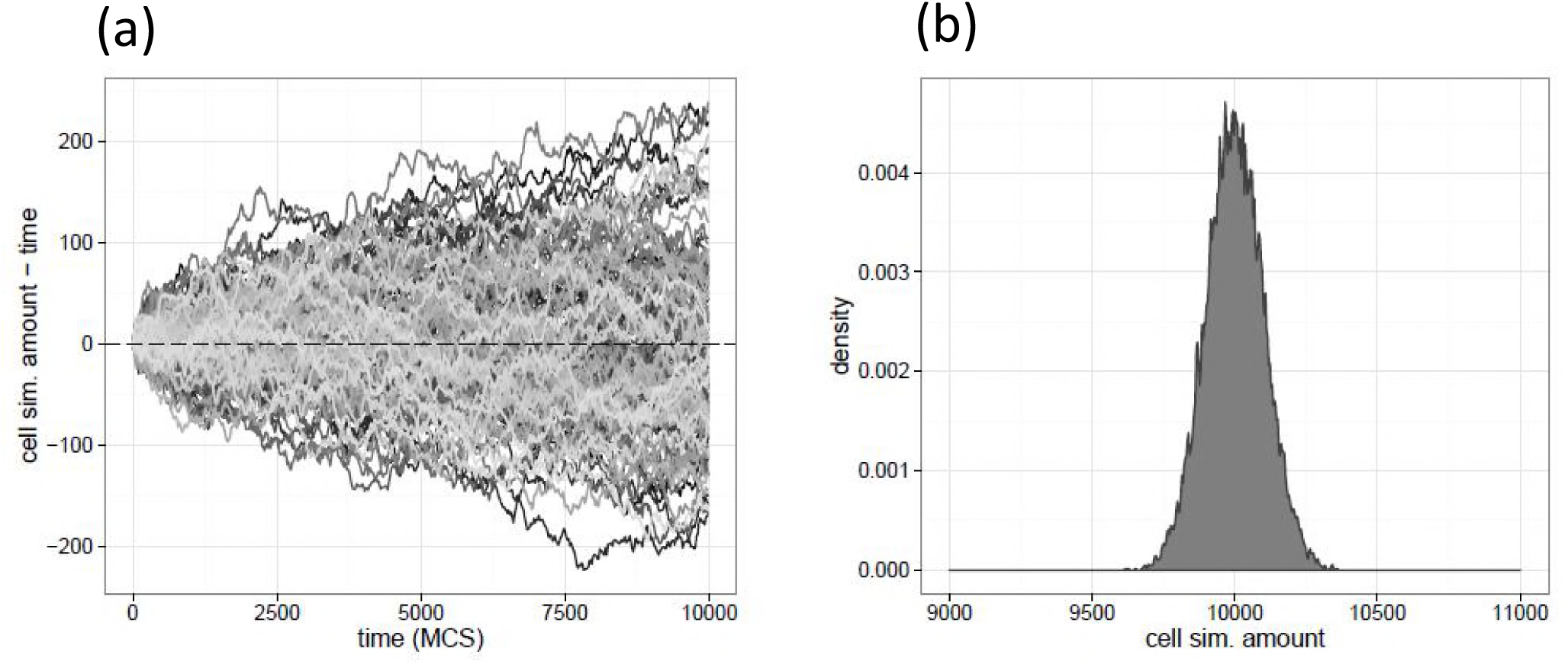
Algorithm Analysis. (a) Selection of trajectories of cells. (b) 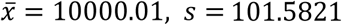.Same parameters as Figure 1, with the additional parameters of *T* = 4, *n*_*C*_ = 213 and *CR* = 2^−13^. Individual cells diverged from a zero mean, and the collective distribution approximated a Gaussian.

## RESULTS AND DISCUSSION

We analyzed the simple substrate-enzyme reaction *S*+ *E* ↔ *C* →*P* + *E*. In this representation, *S* is the substrate, *E* is the enzyme, *C* is a substrate-enzyme complex, and *P* is the product. We will additionally assume that phase-separation occurs in the product, *P*. In our model, *S, E, C*, and *P*, will each occupy one cell with no possibility of overlap, and diffuse with some probability *P*_*dif* *SP*_ for *S* and *P*, and *P*_*dif* *EC*_ for *E* and *C* via a random-walk into the four cardinal directions. Diffusing outside of the lattice removes all particles except for *E* and *C*, which cannot diffuse out. As *S* and *E* attempt to diffuse into each other, both are removed and the site of attempted diffusion is replaced by *C* with the probability of a binding reaction occurring, *P*_*r*_. Failure to bind leaves both *S* and *E* stationary. At each time after attempted diffusion, *C* attempts to react with itself, either “unbinding”’ into *S* and *E* or “reacting” into *P* and *E*. This only occurs if an empty neighboring cell exists, with multiple empty neighboring cells requiring one to be chosen at random. The reaction *C* → *S* + *E* occurs with probability *P*_*CS*_, and the reaction *C* → *P* + *E* occurs with probability *C* → *P* + *E*. Phase-separation in the product is modeled as in [15], whereby the probability of diffusion of *P* is modified by a multiplicative factor 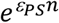, where *ε*_*PS*_.<.0, and *n* is the number of adjacent *P* particles. Thus, a *P* particle with no neighboring *P* particles diffuses with probability *P*_*dif* *SP*_, with one neighboring *P* particle diffusing with probability 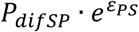, with two neighboring *P* particles diffusing with probability *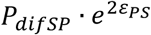.*, with three neighboring *P* particles diffusing with probability *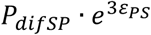*, and with four neighboring *P* particles not being able to diffuse since there are no free adjacent cells to which it is possible to diffuse.

The simulated square *L* × *L* lattice starts with [[*E*]*L*^*2*^].randomly distributed enzymes *E*.Particles *S* enter the lattice, with probability *P*_*S*_._*in*_. when a perimeter cell is simulated (with two independent checks for corners to simulate two possible directions of diffusion). If a particle occupies a perimeter cell, no such check is performed. Let *p*_*dif*_._*SP*_ = 1, *p*_*dif*_._EC_ = 0.1, *P*_*r*_ = 0.01,*p*_*cs*_ = 0.02, *P*_*CP*_ = 0.04, *PS*_.*in*_ = *0.02*5, [*E*] = 0.005. Two values for *ε*_*PS*_.are *ε*_*PS*_ = −3.and *ε*_*PS*_ = –1.5. Both the previous naive and the current cover-based parallel algorithm were implemented in the C programming language [19], with the p-threads library [20] used for parallel computation, and the xoroshiro128+ algorithm used for random number generation. Some calculations were performed with GNUParallel [21]. We analyzed the algorithm in relation to the naive Monte-Carlo method for various *T, n_C_* and *CR*. One quantifiable measure of accuracy is the steady-state of the product, since it depends on the accuracy of the diffusion process.

Figure 6 shows a comparison of the naive simulation for various values of *n*_*C*_, *T*, and *CR*. For *T* = 2 and *T* = 4, it takes very few covers (memory-wise) in order to reach a good level of accuracy. Moreover, there is virtually no difference between the two, and, thus, possibly for larger values of *T*. Additionally, we see that as desired the larger *n*_*C*_, the more accurate the algorithm. As expected, for *CR*, we see that a smaller number is better, with all collisions allowed *CR* = 2^0^ being the worst, and almost no collisions allowed *CR* = 2^−14^ being the best. We can derive an upper bound for the number of collisions expected from a cover, *E*_*c*_ = [*L*^2^ / *CR*]. Thus, per interval, there are *n*_*C*_^*n*^?*E*_*c*_ discrete points of error. As the number *n*_*C*_^*n*^ is completely independent of *n*_*C*_, it should, therefore, follow that for a sufficiently large value of *n*_*C*_, *CR* is the only parameter that determines the accuracy. A choice of a large *n*_*C*_ and small *CR*, therefore, is a good candidate for any given simulation; however, the optimal choice is likely to be problem-based. In order to measure the potential speedup of our algorithm with respect to the naïve variant, we have chosen parameters that are very close to correct values based on our previous analyses. We analyzed the ‘small’; case of *L* = 200, the ‘medium’; case of *L* = 400 (four times larger than ‘small’;) and the ‘large’; case of *L* = 1600 (sixteen times larger than ‘medium’;); we also chose ‘loose’ parameters of *n*_*C*_ = 2^10^ & *CR* = 2^−9^ and ‘strict’; parameters *n*_*C*_ = 2^13^ & *CR* = 2^−13^. In addition, we chose a small constant of phase-separation, *ε*_*PS*_, in order to more quickly reach a steady-state.

**Fig 6.**
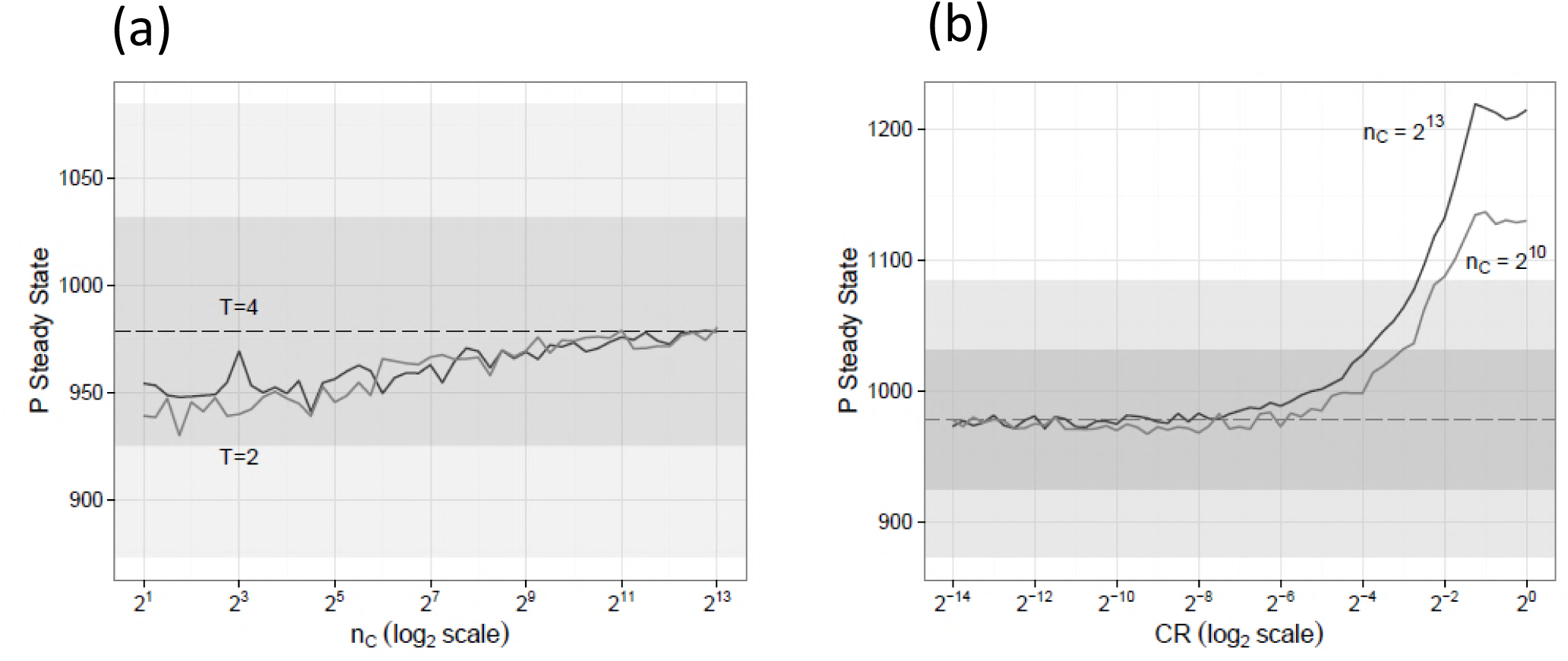
Choosing n_C_, CR, and algorithm accuracy. (a) For *CR* = 2^−9^. (b) For *T* = 4. For *L* = 200, time_MCS_ = 1000000, ΔMCS = 50, and ε_PS_ = –1.5. The dashed line represents the naive algorithm steady-state of *P* with the area around it representing one and two standard deviations. All simulations, including that of the naive model are an averaging of 10 independent runs.

Figure 7 shows the relative speedups of our algorithm compared to the naive variant. We see that for a ‘small’; problem size (*L* = 200), the inefficiency of many threads rapidly manifests itself, which is especially evident for ‘strict’ parameters for which our algorithm is slower than the naive for *T* = 12, 14, 16. This is because for a small enough lattice, we spend more time waiting on a thread barrier than on simulation. For the ‘medium’; problem size, the problem is a lot less evident, as for both ‘loose’; and ‘strict’; parameters the behavior is very similar, but for *T* = 12, a crucial number of threads/cores for the CPU, the ‘loose’; parameter is optimal, but for the ‘strict’; parameter it is not. For the ‘large’; problem size, the behavior is almost exactly the same, except for a slight slowdown for the ‘strict’; parameter set. It is evident that simply utilizing more threads does not speed up the algorithm, and that there is a delicate balance between the problem size (*L*), the initial number of covers (*n*_*C*_), the allowed ratio of collisions (*CR*), and the number of threads (*T*).

**Fig 7.**
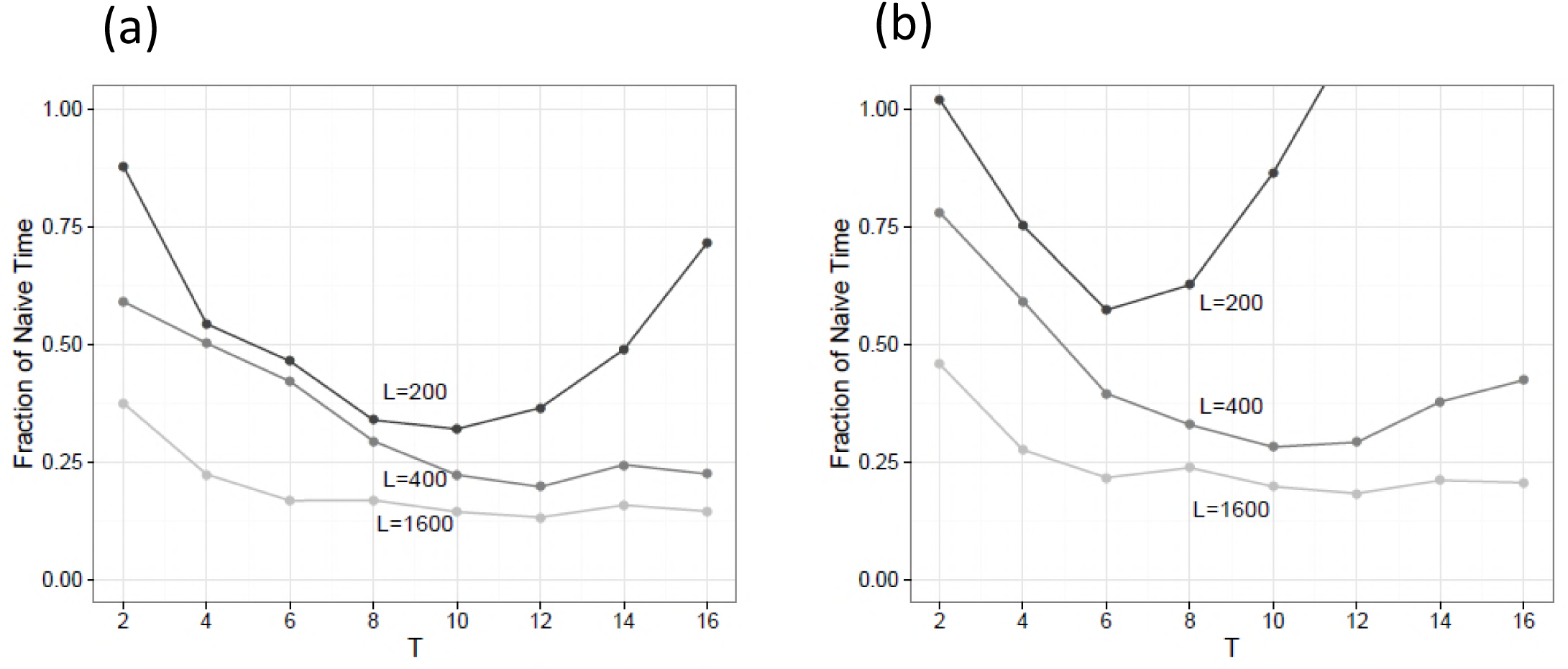
Timing Analysis. (a) For *n*_*C*_ = 2^10^ and *CR* =2^−9^, ‘loose’;. (b) For *n*_*C*_ = 2^13^ and CR = 2^−13^, strict. For time_MCS_ = 250000, ΔMCS = 1000, and ε_PS_ = –1.5. Analysis of the relative speed-up of our algorithm with respect to the naive Monte-Carlo algorithm. On the scale, 1 indicates the time that the naive Monte-Carlo algorithm took to finish the simulation. Obtained on a machine with 2 Intel Xeon E5-2630 processor of 6 cores (2 threads per core) each, which is in accord with the data that indicate that 8 threads of execution were worse than 6 for L = 1600, as 6 is the optimal number for everything to be cached on one physical CPU for this particular setup.

For the substrate-enzyme reaction with phase-separation, a sufficiently large amount of simulation time is required to reach a steady-state. In our simulation of 10^7^ MCS with a more realistic *ε*_*PS*_ = −3 (Fig 8), a near steady-state of the product was achieved after about 2.5 ×10^6^ MCS. However, judging by the empty region in the center of the phase-separated product, even 10^7^ steps are not sufficient to reach a true steady-state. The initial wave of *S* influx is fairly rapid and saturates the lattice quite quickly; thus, the bottleneck seems to be the behavior of the phase-separated particles.

**Fig 8.**
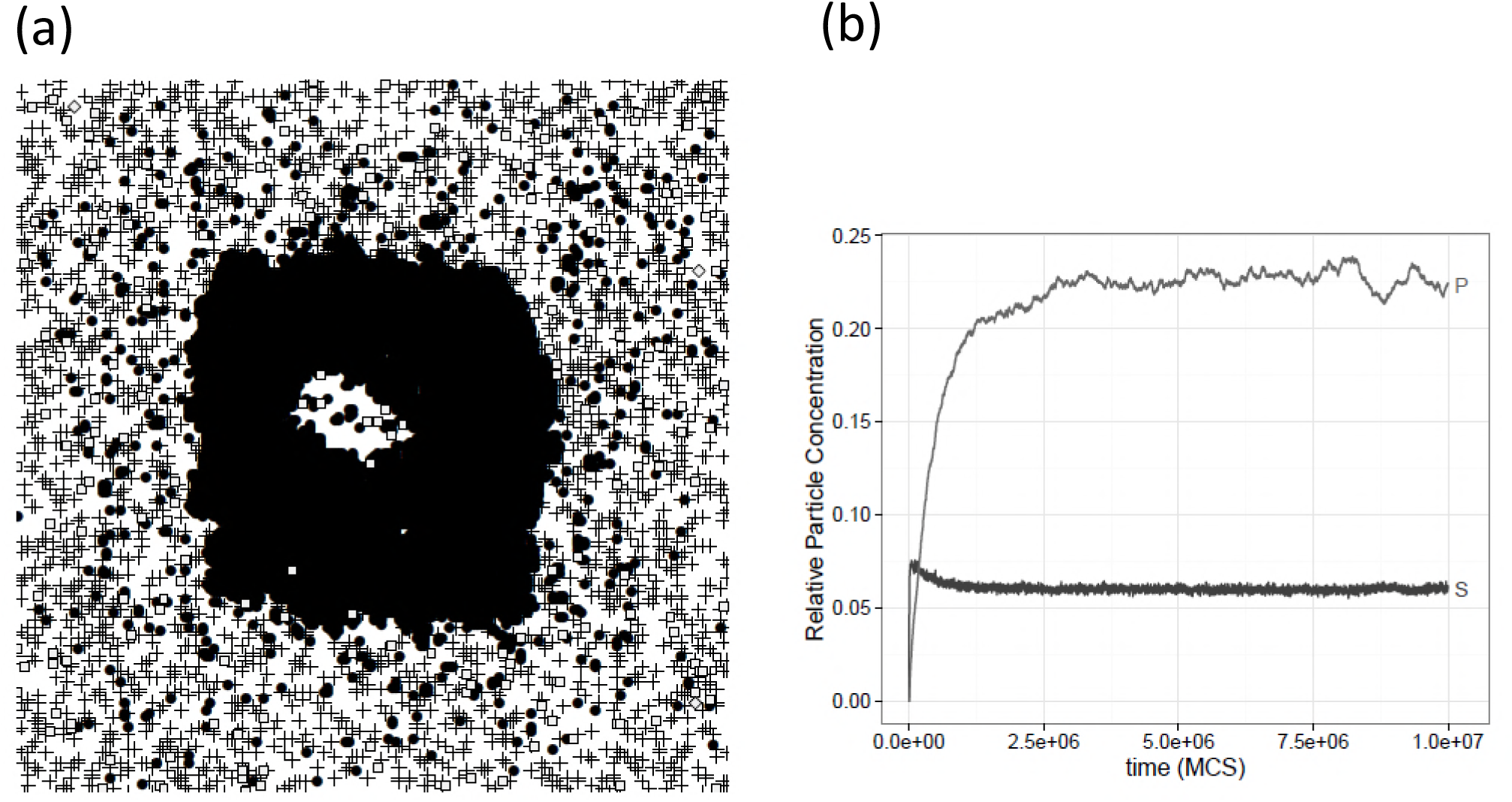
Sample output of the substrate-enzyme model by the parallel algorithm. For *L* = 200,ε_PS_ = −3, time_MCS_ = 10^7^, ΔMCS = 500, *T* = 2, C_n_ = 2^10^, and CR = 2^−9^. The visual image on the right is a spatial representation of the final state of the lattice simulation. The graph on the right is a representation of the state space with respect to time. Enzymes and substrate-enzyme complexes are too small to accurately represent. Even after 10^7^ MCS the simulation had not quite reached a steady-state because of phase separation.

## CONCLUSION

The algorithm that we introduce is both fast and safe for large-scale fractal kinetic simulations with minimal deviation from the naive approach. The speed-up allows us to explore aspects of complex crowded medium interactions such as phase-separation found in various substrate-enzyme reactions. Future work will involve studying how other reactions behave with the addition of phase-separation and a more rigorous examination of the substrate-enzyme reaction.

## AUTHOR CONTRIBUTIONS

A.A.P. and A.A.S. designed research. A.A.P. developed the parallel algorithm and performed numerical experiments. P.P.K. consulted on simulations. A.A.P., S.-C.L., P.P.K., J.D.G., A.A.S. analyzed data. A.A.P. wrote the manuscript. S.-C.L., J.D.G. and A.A.S. edited and reviewed the article.

## ACKNOWLEDGEMENTS

This work was supported in part by grants from the NIH (R01CA172820, R01CA129544 to J.D.G.).

